# Global Land Use Implications of Dietary Trends: A Tragedy of the Commons

**DOI:** 10.1101/195396

**Authors:** Sarah Rizvi, Chris Pagnutti, Chris T. Bauch, Madhur Anand

## Abstract

Global food security and agricultural land management represent two urgent and intimately related challenges that humans must face in the coming decades. Here, we quantify the changes in the global agricultural land footprint if the world were to adhere to the dietary guidelines put forth by the United States Department of Agriculture (USDA), while accounting for the land use change incurred by import/export required to meet those guidelines. We analyze data at country, continent, and global levels. USDA guidelines are viewed as an improvement on the current land-intensive diet of the average American, but despite this our results show that global adherence to USDA guidelines would require up to 1 gigahectare of additional agricultural land--roughly the size of Canada. The results also show a strong divide between Eastern and Western hemispheres. Because countries increasingly import most of their food, meeting USDA guidelines could cause a Tragedy of the Commons, where self-interested actors race to over-exploit the shared resource of global agricultural lands. National dietary guidelines and practices thus need to be coordinated internationally in order to spare our remaining natural lands, in much the same way that countries are coordinating greenhouse gas emissions.

## Introduction

Increasing pressures on land and other natural resources is largely attributed to the increase in sdemand for agricultural products. Global production of food uses approximately 38% of the land on Earth [1]. The agricultural sector is extremely resource-intensive and continues to transform itself as populations grow. An estimated 62% of the remaining global land surface is either unsuitable for cultivation on account of soil, climate topography, or urban development (30%) or is covered in natural land states like forests (32%), so very little land is available for agricultural expansion that does not destroy native land states. Hence, more efficient agricultural production is urgently needed [2]. Currently, approximately 12% of the world remains undernourished [1]. According to estimates from the Food and Agriculture Organization of the United Nations (FAO), the world will need to produce 70% more food by 2050 to meet increased demand [2]. For these reasons, it has become important now more than ever to find effective ways to sustain global agricultural production at healthy and equitable levels.

Diet is an important factor in achieving sustainability in agriculture and equal resource distribution. Food consumption patterns vary widely between countries and cultures. As shown by the FAO, average caloric intake in least developed, developing, and industrialised countries varies widely; 2,120, 2,640, and 3,430 kcal per person per day, respectively [3,4]. However, in many developing countries the average intake is lower than 2,120 kcal per person, resulting in undernourishment [2]. The United States Department of Agriculture (USDA) released *The Dietary Guidelines for Americans, 2010* to promote a healthy diet low in calories and saturated fats. The dietary guidelines are divided by food groups and daily caloric intake levels depending on age, sex, and physiological status [5]. Currently, agricultural outputs and dietary practice— both within the United States and many other rich countries—do not mee those guidelines and favour more land-intensive and calorie-rich diets, such as high levels of meat consumption [6,7]. The global food system is at a point of change where a thorough understanding of the relationship between food consumption patterns, agricultural production and distribution is required to improve the overall sustainability of the system [8].

In this paper we address the questions: If every country were to adhere to the USDA guidelines, what are the implications for agricultural land expansion worldwide? Is there enough land to satisfy the guidelines under current agricultural practice? The global landscape would presumably be greatly altered by the required reallocation of resources. Comparing the recommended food group servings (Table 1) to the food balance sheets reported by the FAO suggests that the average diet in most countries does not match the USDA guidelines. As we will show, to meet these dietary targets, many countries would need to reduce the amount of land they use for producing certain food groups and greatly increase land used for others. Our study determines how land use in the world would change if consumers were to adopt the USDA dietary guidelines for each food group listed in Table 1. In general, to meet the dietary guidelines, we allow that imports may be increased, exports may be changed to domestic production, and domestic production may be expanded where possible. We also discuss how international trade in agricultural products combined with growing land use requirements can create conditions for a Tragedy of the Commons that will require international coordination to avert.

**Table 1.** Daily recommended caloric intake of each food group as outlined by the United States Department of Agriculture Food Guide. Table adapted from the USDA *Dietary Guidelines for Americans 2010* [5]. Food groups are divided into 6 categories with servings determined by caloric levels. The caloric levels are assigned based on sex, physiological status and age.

| Food Group | Daily Calorie Level |  |  |  |  |  |  |  |  |  |  |  |
| --- | --- | --- | --- | --- | --- | --- | --- | --- | --- | --- | --- | --- |
|  | 1000 | 1200 | 1400 | 1600 | 1800<br>(Servings) | 2000 | 2200 | 2400 | 2600 | 2800 | 3000 | 3200 |
| Fruit (Cups) | 1.0 | 1.0 | 1.5 | 1.5 | 1.5 | 2.0 | 2.0 | 2.0 | 2.0 | 2.5 | 2.5 | 2.5 |
| Vegetables (Cups) | 1.0 | 1.5 | 1.5 | 2.0 | 2.5 | 2.5 | 3.0 | 3.0 | 3.5 | 3.5 | 4.0 | 4.0 |
| Grains | 3.0 | 4.0 | 5.0 | 5.0 | 6.0 | 6.0 | 7.0 | 8.0 | 9.0 | 10.0 | 10.0 | 10.0 |
| Whole-grain portion (oz-eq) | 1.5 | 2.0 | 2.5 | 3.0 | 3.0 | 3.0 | 3.5 | 4.0 | 4.5 | 5.0 | 5.0 | 5.0 |
| Meat and Beans (oz-eq) | 2.0 | 3.0 | 4.0 | 5.0 | 5.0 | 5.5 | 6.0 | 6.5 | 6.5 | 7.0 | 7.0 | 7.0 |
| Milk (cups) | 2.0 | 2.0 | 2.0 | 3.0 | 3.0 | 3.0 | 3.0 | 3.0 | 3.0 | 3.0 | 3.0 | 3.0 |
| Oils (tsp) | 3.0 | 4.0 | 4.0 | 5.0 | 5.0 | 6.0 | 6.0 | 7.0 | 8.0 | 8.0 | 10.0 | 10.0 |
| Discretionary calorie allowance | 165 | 171 | 171 | 132 | 195 | 267 | 290 | 362 | 410 | 426 | 512 | 648 |

## Results

We used FAOSTAT data to convert the USDA *Dietary Guidelines for Americans, 2010* to land area required for the guideline diet at the level of country, continent, and world (see Methods for details) [1]. We wished to estimate a conservative lower bound on the amount of land needed to meet the guidelines, if countries were to switch to the USDA guidelines in 2010. Hence, instead of relying on model-based projections, we used historical FAOSTAT country-level data and estimated the amount of land required for the guideline diet, given the observed (lower) historical population sizes and agricultural activity until 2010. Hence, the resulting data point for each year represents the amount of land spared or required in that year, if the given country had been adhering to the USDA guidelines. Although we generated these estimates for 1960 to 2010, the values for 2010 are most relevant to the current situation and generate a lower bound for possible future land requirements and we focus on the 2010 estimates in our Results. We computed both land required under domestic production as well as “displaced” land required—food consumed by a country that was grown on land in another country.

## Global analysis

On a global scale it is apparent that certain food groups are driving changes in agriculture. Looking at these trends we see that overall, if the world were to alter their food consumption to meet the USDA *Dietary Guidelines for Americans, 2010*, there would need to be a dramatic and unsustainable increase in agricultural lands (Figure 1).

**Figure 1.**
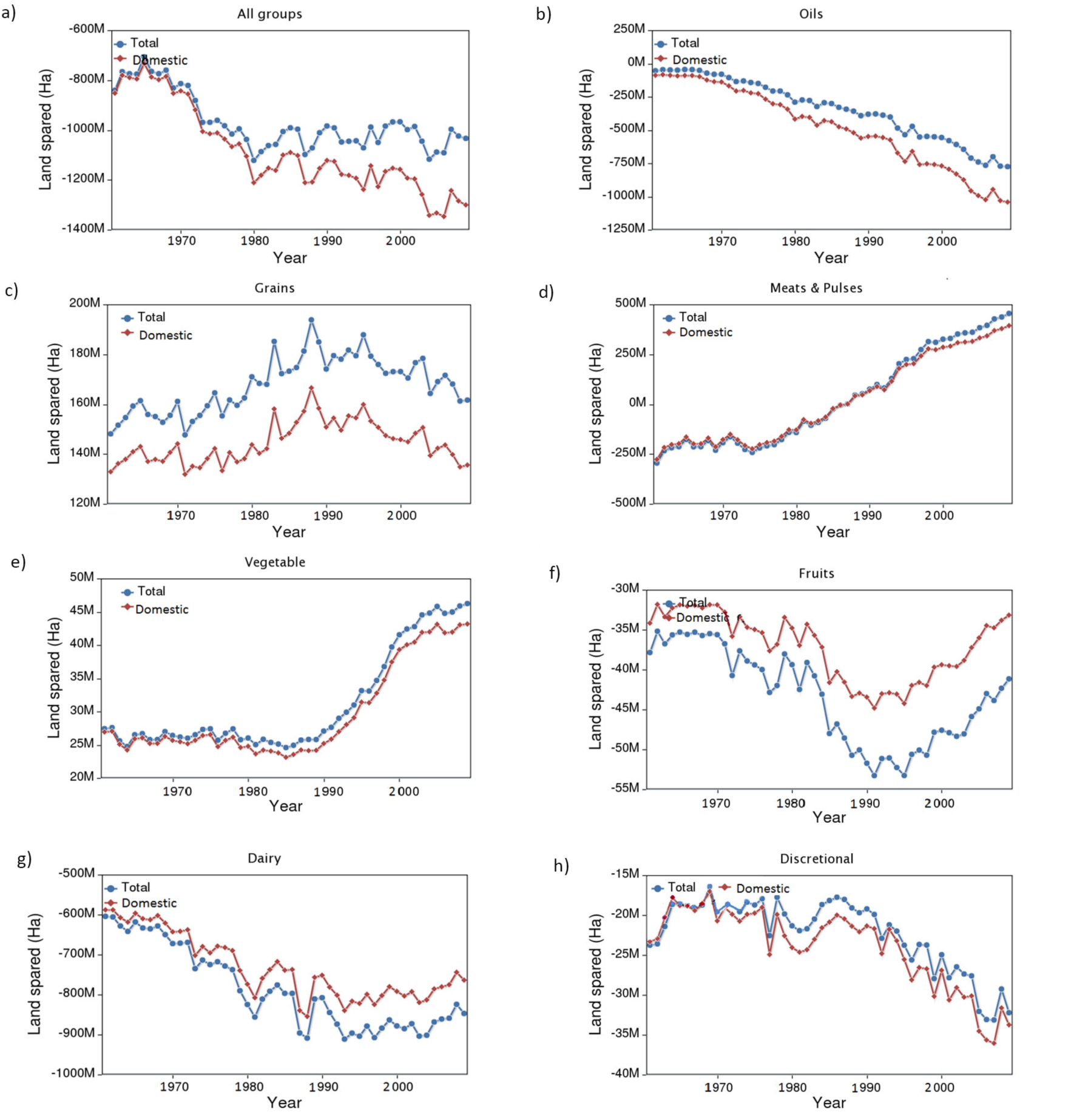
There is not enough land in the world to allow everyone to eat a USDA guideline diet. Plot shows amount of land spared or required to meet the USDA *Dietary Guidelines for Americans 2010*, by year for (a) all food groups, and for (b) oils, (c) grains, (d) meat and pulses, (e) vegetables, (f) fruits, (g) dairy, and (h) discretional. Red depicts the amount of land spared or required based only on domestic production while the blue line combines domestic land and displaced land (land use a country generates elsewhere by relying on food imports) to depict a total amount of land spared or required. A negative amount of land spared means a land deficit: more land will be needed to produce the amount of food required by the USDA guidelines. The gap between domestic and total land spared for all groups is nonzero due to discrepancies in the FAO dataset; the two curves should match one another.

Overall, for the world to meet the guidelines, additional land is required for fruits, dairy and oils and discretional products (Figure 1). In contrast, significant amounts of land could be spared in the meat, vegetables and grain sectors. This trend is common to most continents except Africa (Supporting Information). However, in total for all food groups, approximately 1 gigahectare (Gha) of additional land is required to meet the guidelines (Figure 1, “all groups”, 2010 data point). 1 Gha of land is roughly the size of Canada and exceeds the amount of fertile land currently available worldwide. Hence, even the current USDA guidelines do not go far enough in terms of setting up a globally sustainable dietary practice, from a land use perspective.

Our analysis also shows temporal trends in land spared or required under the guidelines (Figure 1). Required land has been steadfastly increasing since 1960 (Figure 1, “all groups”) due to increasing global populations.

## Analysis by continent

The challenges of providing stable access to adequate food are exacerbated by inequitable dietary patterns of over- and under-consumption between countries and continents. Some of these issues become apparent when we analyze data at the continental level, at which notable common trends in consumption patterns and the associated land requirements emerge. For instance, North America and the European Union displace more land than any other continents, due to food imports (Figure 2). If North and South America shifted to USDA guidelines, they would spare a moderate amount of land from changing to a less land-intensive diet (based on the 2010 estimates). In contrast, if Africa, Eastern Europe, the European Union and Oceania shifted to USDA guidelines, a very large land deficit would develop. The impact of Asia shifting to USDA guidelines would be almost neutral, although the historical trend suggests this will not be the case in the near future. The fact that the European Union (a relatively wealthy part of the world where caloric intake is general adequate as it currently stands) would cause a land deficit by shifting to the USDA guidelines suggests that the USDA guidelines are unsustainable when it comes to land-intensive food groups like meat.

**Figure 2.**
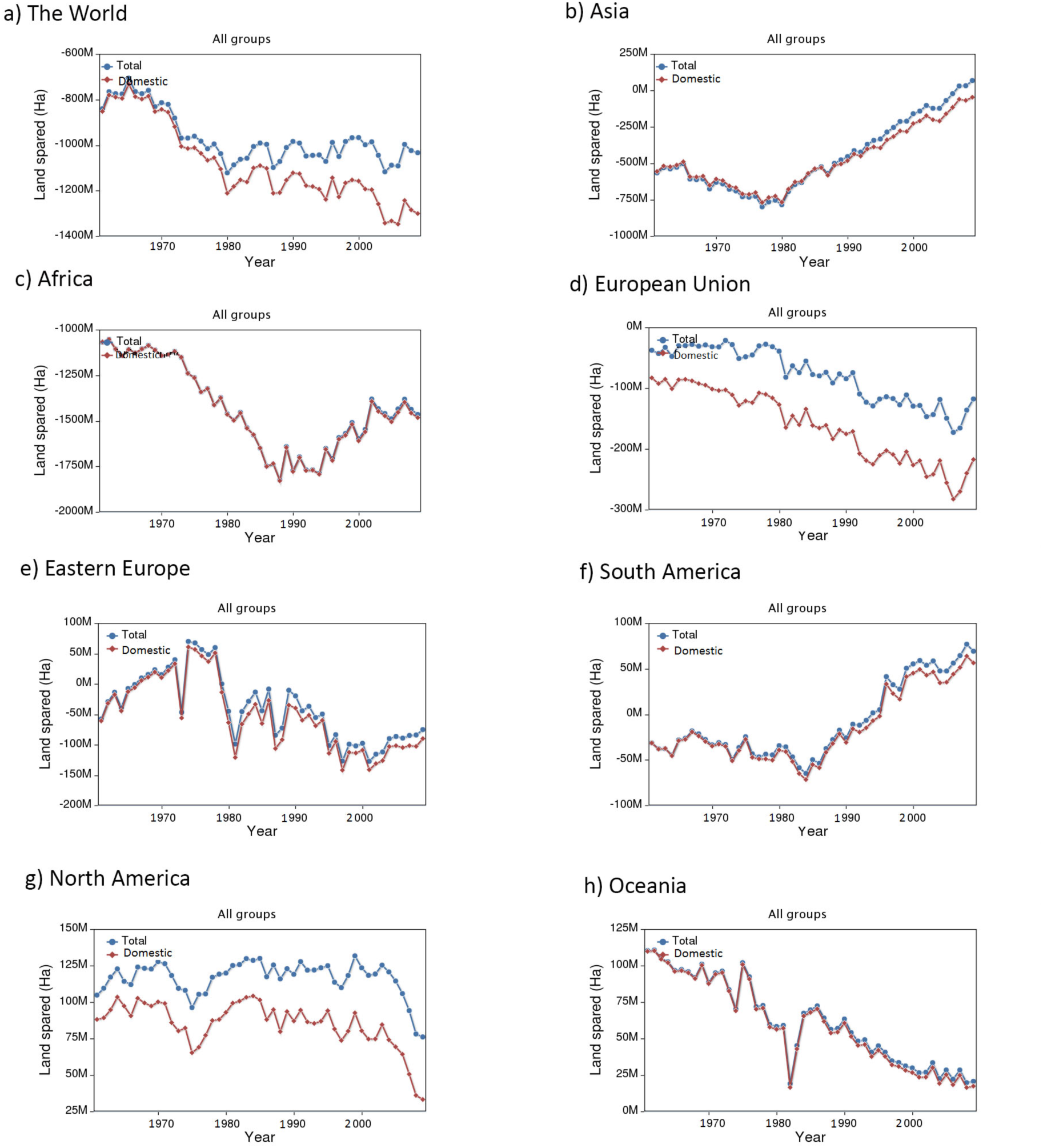
Continents differ widely in land spared or required under USDA guideline diet. Plots show change in agricultural land area required to meet the *Dietary Guidelines* in each continent, by year for (a) the world, (b) Asia, (c) Africa, (d) European Union, (e) Eastern Europe, (f) South America, (g) North America and (h) Oceania. Red depicts the amount of land spared or required based only on domestic production while the blue line combines domestic land and displaced land (land use a country generates elsewhere by relying on food imports) to depict a total amount of land spared or required. A negative amount of land spared means a land deficit: more land will be needed to produce the amount of food required by the USDA guidelines.

For most decades in the Asia dataset, Asia would have caused a net land deficit by shifting to the USDA guidelines, since it was (and remains) a relatively under-nourished part of the world (Figure 2b). An inflection point appears in the Asian dataset in 1980, when countries like China and India began liberalizing their economies. In the most recent years, Asia is estimated to cause a small amount of land sparing due to economic growth, should it switch to the USDA guidelines. Most notably are increases in land use for meat and grains as Asia slowly begins to adopt a more westernized diet (Supplementary Information). This suggests that while Asia has increased land use rapidly, equity in resource distribution at the sub-continental level is imbalanced. For instance, one third of Indians are undernourished and continue to live in food insecurity [2]. Inequities in global trading and extension services as well as poor infrastructure trap populations in Asia in poverty. However, future improvements towards equal land use change may better harness agricultural yields to align the Asian diet with those of wealthier and more sustainable areas of the world, such as the European Union.

Africa would require more land to meet the USDA guidelines than any other region. In fact, most of the additional land required to meet the guidelines globally would be the result of dietary shifts in Africa (using 2010 as the reference point). This is not surprising because undernourishment is widespread in Africa [12]. However, an inflection point, probably corresponding to growth in some African economies, occurs in 1990 (Figure 2c). Almost all of the additional land required to meet the guidelines would be the result of increased dairy consumption (Supporting Information). Although the extra land required to meet the guidelines in Africa is impossibly large (more land is needed than what is available), Africa also stands the most to gain with respect to growing agricultural yields [13]. Thus, although it is not currently possible to bring the African diet in line with that of the USA or much less the comparatively land-sparing diet of the European Union without a net growth in agricultural lands, future improvements in agricultural practices in Africa may help to close the gap.

The European Union would also require a significant amount of land to meet the USDA guidelines. Almost all of the additional land needed would be the result of increased dairy and fruit land use, a trend common to most of Europe (Supporting Information). We note that displaced land (from buying food imports) contributes strongly to European Union land use, and exceeds displaced land use in North America (Figure 2d). Interestingly, the land requirements for the European Union indicate the need for more displaced lands than domestic land. This suggests that an American diet is unsustainable from a land use perspective, domestically speaking.

Land use in Eastern Europe has fluctuated significantly over time (Figure 2e). After the late 1980s, a land use deficit developed in the Eastern Europe dataset, and has largely persisted in recent years. Therefore, in order to meet the USDA guidelines, Eastern Europe would require a large amount of new land.

North America can spare a significant amount of land, should the USDA guidelines be followed. The sparing stems largely from meat, grain and vegetable land use (Supporting Information) [14]. Land use for meat is greater in North America than any other continent, and as a result, land use displacement in North America is also significant (Figure 2g).

South America can also spare a significant amount of land in order to meet the USDA guidelines, mostly from land sparing due to meat and grains, followed by vegetables and discretional products. South America shows a steady increase in land use since 1984 (Figure 2f). This trend is overwhelmingly due to rapid increases in land use for meat. Thus, reducing meat consumption in South America shows strong potential for sparing land (Supporting Information). According to United Nations Environment (UNEP), ranches alone accounted for an estimated 70% of deforestation in Brazil in 2007, where ranches covered approximately 8.4 million hectares [2]. Finally, Oceania can spare a small amount of land if the guidelines are met, primarily from meat, grains and vegetables (Figure 2h, Supporting Information).

## World Map

Using the FAOSTAT data and USDA guidelines [1,5] we also created a world map (see Methods) showing land spared or required for shifting to the USDA guidelines for each country as of 2009 (Figure 3). The countries that can spare the most land if dietary guidelines are met are the United States of America, Brazil and Australia. In contrast, the countries that require the most land to meet the guidelines are Mozambique, Saudi Arabia, and India. The country level map also shows a strong hemispheric divide, with the Western hemisphere able to spare significant amounts of land by moving to the USDA guidelines (due largely to higher consumption of meat, and grain grown to feed livestock), while the Eastern hemisphere would require net new land in order to have a diet similar to the USDA guidelines.

**Figure 3.**
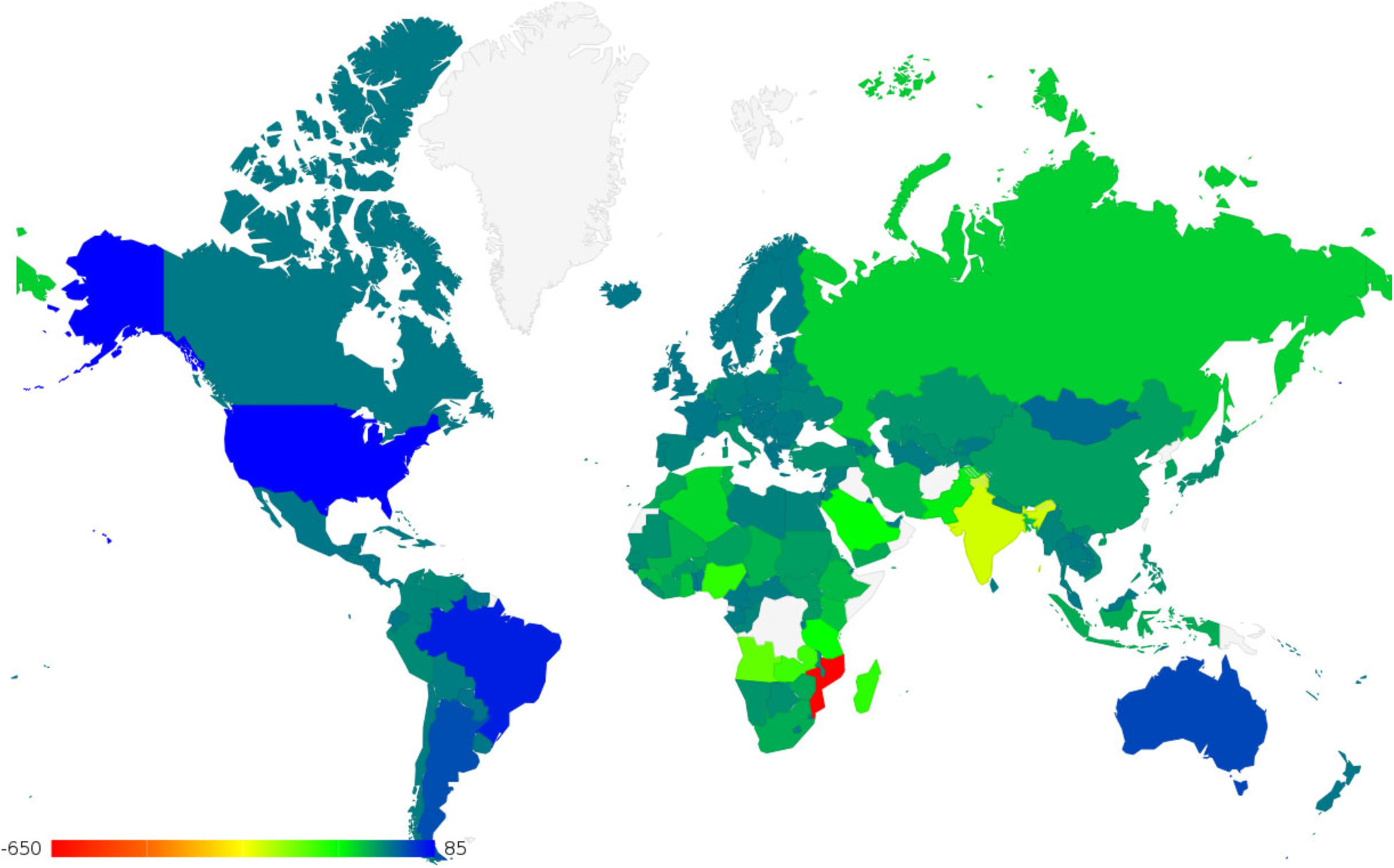
A western/eastern hemispheric divide in land spared versus land required by a USDA guideline diet. Land spared or required in 2009 on a global scale. The scale depicts negative surplus as a deficit beginning in red, meaning that the country would need more food and more land to follow the guidelines. The scale progresses to blue depicting a positive surplus, meaning the country would need less food and less land to follow the guidelines. The map was created by the authors using the Google Maps API (https://developers.google.com/maps/ with Apache License Version 2.0) based on the FAOSTAT dataset [1].

## Discussion

The USDA *Dietary Guidelines for Americans, 2010* already promotes a more balanced diet than that practiced by most Americans and to which many individuals in countries with rapidly expanding economics aspire. However, our analysis shows that there is not enough land for the world to adhere to the USDA guidelines under current agricultural practices. A staggering 1 gigahectare of fertile land—roughly the size of Canada—would be required. Moreover, our estimate is arguably conservative since we relied upon recent historical data rather than attempting to project forward using population and economic models. However, it also bears pointing out that we thereby neglect possible future increases in agricultural yield in continents like Africa. Despite this, the required amount of land is massive enough to raise concerns about our ability to feed everyone in the world in an equitable and a sustainable way. New technologies such as lab-grown meat have been invoked as part of the solution to this crisis, but new technologies seem perpetually 5-10 years away and have unknown prospects for adoption when they finally do arrive. We conclude that a change in dietary trends is a necessary part of the solution to this problem.

The analysis was also broken down by continent and country. The western hemisphere (North America and South America) and Oceania could spare significant amounts of land if they moved to the less meat-intensive (and consequently, grain-intensive) diet suggested in the USDA guidelines. In contrast, eastern hemispheric continents (Africa, European Union and Asia) would require a significant expansion of agricultural lands to support a USDA guideline diet. Recent dietary trends in Africa and Asia (Figure 2) show movement toward these guidelines, as reflected in other research on evolving diets in these regions [15]. China, India, and Saudi Arabia have changed most drastically in recent years with an increase in agricultural land use. Pakistan, along with India, is responding to growing consumer demand for more western diets by increasing beef production [16]. Of particular interest in Asia is China, which is rapidly increasing production in several sectors, largely contributing to Asia’s rapid agricultural growth rate (Supporting Information) [15]. Humans will have to deal with growing inequities as growing land use for meat consumption by richer individuals and richer countries causes rising food costs for staples such as pulses and grains and thus harms the poor and under-nourished remainder [17,18].

Future research could study the impact of real or potential dietary shifts on greenhouse gas emissions. Global agricultural production accounts for nearly 30% of total greenhouse gas (GHG) emissions [18]. Livestock alone are responsible for 18% of GHG emissions, which is higher than the share of GHG emissions from transportation [16]. Hence, a shift to less meat consumption would also reduce GHG emissions. Other topics for future research include the effects of food lost during storage and transportation and (more importantly) food lost through waste and disposal. Food loss is significant around the world, thus reducing food loss could also help spare land.

Finally, these results highlight how dietary shifts are creating a “Tragedy of the Commons” in global land use. In 1968 Garrett Hardin coined this term to describe common resource systems where self-interested individuals, in the absence of social norms or regulations, over-exploit a common resource and thereby generate a result that harms the common good [19-21]. The Tragedy of the Commons has since been used as a conceptual framework for phenomena such as anthropogenic climate change and fishery collapse. Because the Earth’s atmosphere is a well-mixed system, one country’s emissions affect the climate of the entire planet. Therefore in the absence of international coordination, any individual country has little incentive to reduce their GHG emissions, if no other countries are also reducing theirs. Historically, most food was produced and consumed in the same country and hence an international Tragedy of the Commons in land use was not possible (although small scale overexploitation of a village’s grazing commons was the origin of the concept, ironically). However, the rapid growth of trade in agricultural products is such that many countries now rely on food imports [22]. Moreover, although any given agricultural plot at any given time is owned by a single entity and not shared, on larger spatial and temporal scales the emerging globally mixing system of “virtually traded lands” [23] represents a *de facto* commons. Thereby it creates conditions similar to those created by GHG emissions or fisheries exploitation. In light of the results of our study that there is not enough land for everyone to eat according to the USDA guidelines, this is creating a global Tragedy of the Commons in land use.

We suggest that global agricultural activity and the corresponding land use challenges should therefore be framed as a Tragedy of the Commons. The implication of this change in framing is that countries should coordinate their formulation of dietary guidelines such that they are based not only on health considerations but also considerations of sustainable global land use and natural ecosystem conservation. Moreover, international coordination should incentivize country-level improvements in dietary habits that result in global land sparing, similar to how countries are beginning to coordinate reductions in their GHG emissions.

## Methods

We chose to use the USDA *Dietary Guidelines for Americans 2010* because the guidelines are comprehensive and well-articulated [5]. Also, as a global power, the United States partially determines global standards.. More developing nations are beginning to adopt a more westernized lifestyle including a diet similar to that expressed in the USDA guidelines, so the study is consistent with ongoing global dietary trends.

We used the FAOSTAT database [1] to compile the food supply quantity for each of the commodity aggregates listed in Table 1 and grouped them according to the major food groups recognized in the USDA MyPyramid model: fruits, vegetables, grains, meat/protein, dairy, oils and discretional [5]. For beverages, oils, sugar, butter and stimulants we converted the processed quantities to equivalent primary quantities (e.g. wine to grapes, beer to barley, butter to milk etc.) using conversion factors given by the FAO [9]. We also computed the import dependency ratio, defined as the ratio of the import quantity to the domestic supply quantity, for each country and each commodity.

Next we took the recommended daily serving sizes of each food group based on the 2000 kcal/day level and converted those to masses using the food balance sheets handbook given by the FAO [10]. For each country we multiplied each of these masses by 365 (days) times the population of the country to get the quantity of each food group that would be required in order for that country to adhere to the USDA guidelines in a year. A country’s surplus of each food group was taken to be the actual food supply for each food group minus the corresponding quantity that would be required to meet the USDA guidelines. A negative surplus is to be interpreted as a deficit, meaning that the country would need more food from that group to follow the guidelines.

For each country the surplus of each food group was divided into two parts: one that was produced within that country (domestic), and one that was produced outside of that country (displaced) according to the import dependency ratio [10], which is defined as the ratio of the import quantity to the domestic supply quantity, and which can be calculated from data in the food balance sheets in the FAOSTAT database. Finally, for the domestic portion the change in agricultural land area within that country that is required to meet the USDA guidelines was taken to be the domestic surplus divided by that country’s combined yield of all commodities in the given food group (Table 2). The change in agricultural land area outside of that country was computed in the same way, but using the displaced surplus and the world average yields. Yield data for crops can be found in the FAOSTAT database. Yield in terms of production per hectare of land used for livestock products was previously calculated [11]. The details of the calculations are described in the python script given in the Supporting Information. Code used for analysis can be downloaded from https://github.com/Pacopag/faolyzer.

**Table 2.** FAO items, codes and the corresponding food group. The codes without parentheses refer to commodity aggregates reported in the food balance sheets, and the codes in parentheses refer to the corresponding items in the “Production” data in the FAOSTAT database [1]. The conversion factors were used to convert quantities to their primary item equivalent (e.g. wine gets converted to equivalent quantity of grapes) [7].

| Commodity | FAO codes | Food group | Conversion |
| --- | --- | --- | --- |
| Fruits | 2919 (1801) | Fruits | - |
| Wine | 2655 (560) | Fruits | 0.7 |
| Vegetables | 2918 (1735) | Vegetables | - |
| Starchy Roots | 2907 (1720) | Vegetables | - |
| Cereals | 2905 (1717) | Grains | - |
| Beer | 2656 (44) | Grains | 4.78 |
| Beverages, Alcoholic | 2658 (1717) | Grains | 0.6 |
| Bovine meat | 2731 (867) | Meat/Protein | - |
| Mutton and Goat meat | 2731 (977,1017) | Meat/Protein | - |
| Pig meat | 2733 (1035) | Meat/Protein | - |
| Poultry meat | 2734 (1058) | Meat/Protein | - |
| Eggs | 2744 (1062) | Meat/Protein | - |
| Oil crops | 2913 (1732) | Meat/Protein | - |
| Treenuts | 2912 (1729) | Meat/Protein | - |
| Pulses | 2911 (1726) | Meat/Protein | - |
| Milk | 2948 (882) | Dairy | - |
| Butter, Ghee | 2740 (886) | Dairy | 0.047 |
| Oils | 2914 (1732) | Oils | 0.2 |
| Sugar crops | 2908 (156,157,161) | Discretionary | - |
| Sugar and Sweeteners | 2909 (156,157,161) | Discretionary | 0.12 |
| Stimulants | 2922 (661,656) | Discretionary | - |

The visualization of our country-level data in Figure 3 was generated using the Google Maps API [24].

## Acknowledgments

This research was supported by an NSERC Discovery Grant to M.A.

## Author Contributions

M.A. conceived of the study. All authors designed the analysis. S.R. and C.P. conducted the analysis. All authors contributed to writing the manuscript.

## Additional Information

The authors have no competing interests to declare.

